# Beyond PAM50: Unsupervised Discovery of Anomalous Subgroups in Breast Cancer

**DOI:** 10.1101/2025.05.11.653361

**Authors:** Tushar Gairola

## Abstract

Breast invasive carcinoma (BRCA) exhibits molecular heterogeneity not fully captured by classifiers like PAM50. I applied an ensemble of four unsupervised anomaly detection algorithms Isolation Forest, One-Class SVM, Local Outlier Factor, and Autoencoder to ∼13,400 gene expression profiles from 1,218 TCGA-BRCA RNA-seq samples, identifying 41 High-Concordance Anomalies (HCAs) consistently flagged by three or more methods. HCAs showed marked downregulation of ∼1,750 genes, strongly enriched for immune-related pathways such as T-cell activation and cytokine signaling, indicating an “immune-cold” phenotype. In contrast, ∼160 upregulated genes were associated with metal ion response, metabolism, and developmental programs. Over half of the HCAs were PAM50_Unknown. Within the Basal-like subtype, a subset of HCAs (HCA-Basal, n=7) exhibited even stronger immune suppression, with 499 additional immune genes downregulated, defining an “ultra-immune-cold” variant. Upregulated genes in HCA-Basal lacked coherent pathway enrichment. While not statistically significant, HCA-Basal cases (n=5 with survival data) showed a trend toward poorer prognosis. These findings reveal a distinct, immune-suppressed BRCA subgroup often missed by current classifiers, with potential relevance for risk assessment and treatment.

## Introduction

Breast invasive carcinoma (BRCA) remains a significant global health burden, representing the most commonly diagnosed cancer and a major contributor to cancer-related deaths among women [Sung et al., 2021]. A hallmark of BRCA is its complex heterogeneity both between and within tumors manifesting as variations in molecular signatures, pathological features, clinical behavior, and treatment responses. This diversity underscores the necessity for precise stratification of patients to enable personalized therapies and enhance clinical outcomes. Over the years, researchers have sought to identify biologically distinct and clinically meaningful subgroups within BRCA.

Traditional classification methods for BRCA relied on histological attributes, including tumor grade and hormone receptor expression (ER, PR, HER2). However, advances in transcriptomic profiling through microarrays and RNA sequencing have enabled the discovery of intrinsic molecular subtypes. Groundbreaking work by Perou, Sørlie, and others established gene expression-based categories Luminal A, Luminal B, HER2-enriched, and Basal like collectively referred to as PAM50 subtypes. These subtypes hold prognostic and therapeutic significance, guiding decisions on endocrine and chemotherapy treatments. Large-scale efforts such as The Cancer Genome Atlas (TCGA) further refined this molecular taxonomy, exposing deeper biological layers [TCGA Network, 2012].

Despite these insights, a notable proportion of BRCA tumors up to 20% remain unclassified by PAM50, termed “PAM50_Unknown” [Akbani et al., 2014]. Additionally, heterogeneity within existing subtypes remains unresolved. For instance, Basal-like tumors show broad variability in outcomes, and even low-risk Luminal A cases may relapse late, indicating missed molecular distinctions. Prior studies have largely focused on optimizing known subtypes, with limited work on identifying tumors that deviate markedly from known profiles.

This study addresses that gap by employing multiple unsupervised anomaly detection algorithms on TCGA-BRCA RNA seq data to identify and characterize high-concordance outlier samples systematically. These “High Concordance Anomalies” (HCAs) are further analyzed for differential expression, pathway enrichment, clinical correlation, and subtype-specific insights, especially within Basal like BRCA. This transcriptomics-focused analysis provides a foundation for uncovering novel tumor subgroups and generating hypotheses for future functional validation

## Materials and Methods

### Data Sources and Cohort Description

I utilized publicly accessible datasets from the TCGA Breast Invasive Carcinoma (TCGA-BRCA) cohort, made available through the [UCSC Xena](https://tcga.xenahubs.net/) Browser.

- **Gene Expression Data:** RNA-Seq expression profiles were retrieved from the “gene expression RNAseq - IlluminaHiSeq” dataset (file: TCGA.BRCA.sampleMap/HiSeqV2, downloaded as TCGA-BRCA_expression_log2norm). These Level 3 RSEM-normalized counts represent log2(norm_count + 1) transformations, produced by the TCGA genome characterization center at UNC, and cover 1,218 primary tumor samples.
- **Clinical and Subtype Data:** Clinical metadata were obtained from the “Phenotypes” dataset (TCGA.BRCA.sampleMap/BRCA_clinicalMatrix, version 2019-12-06). The analysis focused on several essential clinical attributes, including the PAM50 molecular subtype classification (PAM50Call_RNAseq), overall survival duration and corresponding event status (OS_Time_nature2012 and OS_event_nature2012), the patient’s age at the time of initial pathological diagnosis, and the pathological stage of the tumor.

Only patients with overlapping records in both gene expression and clinical datasets were included, resulting in a final analysis cohort of 1,218 samples.

### Preprocessing and Feature Selection

The original gene expression matrix (genes × samples) was transposed to samples × genes. To ensure consistency, we intersected patient IDs across expression and clinical data. The following preprocessing steps were then applied:

1. Low Expression Filtering: Genes with a mean log2(norm_count + 1) < 0.5 across all samples were excluded.
2. Low Variance Filtering: Genes within the lowest 25th percentile of variance were removed.
3. This resulted in a high-quality expression matrix (X_processed) containing 13,407 genes and 1,218 samples, used for all downstream analyses.

### Anomaly Detection Pipeline

#### Feature Scaling

- For most models, X_processed was standardized (zero mean, unit variance per gene) using StandardScaler from scikit-learn.
- For the Autoencoder, I applied MinMax scaling to rescale values between 0 and 1.

#### Anomaly Detection Models

I implemented four unsupervised anomaly detection (AD) algorithms, each initialized with a contamination rate of 0.05:

- Isolation Forest (IF): sklearn.ensemble.IsolationForest with n_estimators=200, max_samples=‘auto’, random_state=42.
- One-Class SVM (OCSVM): sklearn.svm.OneClassSVM using RBF kernel, gamma=‘auto’, nu=0.05.
- Local Outlier Factor (LOF): sklearn.neighbors.LocalOutlierFactor with n_neighbors=30, contamination=0.05, novelty=False.
- Autoencoder (AE): A deep feedforward model implemented in TensorFlow/Keras with:
  - Architecture: Input layer (13,407), encoding layers (670 → 335 → 167), mirrored decoder layers.
  - Activation: ReLU for all layers, Sigmoid for output.
  - Regularization: L1/L2 (1e-5), Dropout (rate 0.1).
  - Optimization: Adam optimizer (lr=0.001), MSE loss, batch size = 32.
  - Training: Early stopping (patience = 15), learning rate reduction on plateau (factor = 0.2, patience = 7).
  - Anomalies were defined as samples with reconstruction errors above the 95th percentile.

#### Concordance Voting

Each model produced binary anomaly predictions (1 = outlier, 0 = normal). A Concordance Vote score (range: 0–4) was calculated per sample. Samples flagged as anomalies by ≥3 models were labeled High-Concordance Anomalies (HCAs). Samples not flagged by any model (0 votes) formed the normal baseline group.

### Differential Gene Expression Analysis (DGE)

We compared gene expression profiles of:

- HCAs vs. 0-vote Normals (global comparison)
- HCA-Basal vs. 0-Vote-Normal-Basal, restricted to PAM50 Basal subtype

For both, i applied Welch’s t-test (via scipy.stats.ttest_ind, equal_var=False) on the original normalized (but not standardized) matrix. Multiple testing corrections used Benjamini-Hochberg FDR (statsmodels.stats.multitest.multipletests). Genes with:

- FDR < 0.05
- |log2 Fold Change| > 1.0

were considered significantly differentially expressed.

### Functional and Pathway Enrichment

Lists of up- and down-regulated genes were submitted to Metascape (https://metascape.org, accessed May 2024), using default “Express Analysis” settings for Homo sapiens. Enrichment analyses spanned:

- GO Biological Processes
- KEGG pathways
- Reactome pathways
- MSigDB Hallmark gene sets

Significance was determined by q-values and enrichment p-values.

### Survival Analysis

Overall Survival (OS) was assessed using Kaplan-Meier estimation and log-rank tests, implemented via the lifelines Python library.

- OS time and status were taken from OS_Time_nature2012 and OS_event_nature2012.
- Samples missing survival data were excluded.
- Key comparisons:
  - HCAs vs. Other Samples (0–2 votes)
  - HCA-Basal vs. Other Basal (0-vote Basal + remaining Basal)

### Software and Visualization

All analyses were conducted in Python 3.11. Core libraries included:

- pandas (v2.x), numpy (v2.x) were used for data handling
- scikit-learn (v1.x) was used for Preprocessing and AD models
- TensorFlow (v2.x) was used for Autoencoder model
- lifelines (v0.x) was used for Survival analysis
- scipy, statsmodels were used for Statistical testing and FDR
- matplotlib, seaborn were used for Visualizations
- umap-learn (v0.5.x) was used for Dimensionality reduction

## Results

### Identification and Visualization of High-Concordance Anomalies in TCGA-BRCA

To explore molecular outliers within the TCGA-BRCA dataset, gene expression profiles from 1,218 tumor samples each containing 13,407 filtered genes were analyzed in conjunction with corresponding clinical data. Four unsupervised anomaly detection (AD) methods were employed: Isolation Forest (IF), Local Outlier Factor (LOF), Autoencoder (AE), and One-Class Support Vector Machine (OCSVM). The first three algorithms each flagged 61 samples as anomalous, while OCSVM identified a broader set of 84 anomalies using a 5% contamination threshold.

A concordance analysis was conducted to enhance the robustness of anomaly detection. This revealed a tiered distribution: 20 samples were flagged by all four models, 21 by three, 30 by two, and 64 by only one. The remaining 1,083 samples unflagged by any method were designated “0-vote normals” and served as a reference group. Samples flagged by at least three methods (41 in total) were classified as “High-Concordance Anomalies” (HCAs), representing robust molecular outliers.

To visualize these findings, UMAP dimensionality reduction was applied, revealing clusters corresponding to PAM50 subtypes. Basal-like and Normal like tumors appeared more distinct, while Luminal A and B tumors showed some overlap. When samples were colored by anomaly concordance votes, HCAs typically appeared at the peripheries or formed distinct clusters, indicating unique expression profiles. These 41 HCAs were highlighted over PAM50 subtype clusters to emphasize their divergence from canonical tumor subtypes.

### Molecular Landscape of High-Concordance Anomalies

Differential gene expression analysis between the 41 HCAs and the 0-vote normals uncovered substantial transcriptomic shifts: 160 genes were significantly up-regulated, and 1,744 were significantly down-regulated (FDR q<0.05, |log2FC|>1). Notably, many of the suppressed genes were immune-related, including **DTHD1, GPR15, CCR4, IL2**, and **CD226**, suggesting an immune-inert tumor phenotype.

Pathway enrichment analysis of the down-regulated genes using Metascape highlighted immune-related GO terms such as “cell activation,” “regulation of cell activation,” and “lymphocyte activation involved in immune response.” KEGG and Reactome analyses reinforced this theme, identifying cytokine-cytokine receptor interactions and interferon signaling pathways as significantly suppressed. Hallmark gene sets like **INTERFERON GAMMA RESPONSE** and **INFLAMMATORY RESPONSE** were also strongly enriched, consolidating the “immune-cold” identity of HCA tumors.

In contrast, up-regulated genes in HCAs were enriched for processes such as metal ion response, energy metabolism, and extracellular matrix organization. Some genes were also linked to neuropeptide signaling and developmental reactivation, hinting at potential dedifferentiation or lineage plasticity. While the immune suppression was dominant, these additional traits may contribute to the distinctive biology of HCAs.

A clustered heatmap of the top differentially expressed genes revealed a clear separation of HCAs, characterized by coordinated down-regulation of immune genes and selective up-regulation of alternative biological programs.

### Clinical Features and Subtype Distribution of HCAs

PAM50 subtype analysis of the 41 HCAs revealed that over half (23/41; ∼56%) were classified as “PAM50 Unknown,” significantly exceeding the ∼20% rate among non-HCA samples. Among HCAs with defined subtypes, Basal-like tumors were somewhat overrepresented (7/41; ∼17%) compared to their prevalence (∼12%) in the rest of the dataset.

Clinical characteristics such as age at diagnosis were similar between HCAs and non-HCAs. Both groups had a mean age of approximately 58 years, and Welch’s t-test yielded a non-significant p-value (0.9448), suggesting no age-related bias in anomaly detection.

### Survival Outcomes

Kaplan-Meier analysis assessed overall survival (OS) in HCAs versus non-HCAs. Among the samples with survival data, 19 HCAs and 914 non HCAs were included. No statistically significant survival difference was found between these groups (log-rank p = 0.2637).

A focused analysis within the Basal-like subtype compared five HCA-Basal samples with survival data to 131 non HCA Basal tumors. While all five HCA-Basal cases experienced a survival event within ∼1,000 days, the small sample size limited statistical power and the difference remained non-significant (log-rank p = 0.1387). Nevertheless, the trend suggested potentially worse prognosis for HCA-Basal tumors.

### Unique Molecular Features of the HCA-Basal Subgroup

To better understand the distinct biology of HCA-Basal tumors, differential expression analysis compared the seven HCA-Basal samples to 113 Basal 0-vote normals. This revealed 15 up-regulated and 499 down-regulated genes (FDR q<0.05, |log2FC|>1). All top-ranked differentially expressed genes were down-regulated, including immune regulators such as **CSF2 (GM-CSF), DPPA4**, and **TUSC5**, suggesting marked suppression of immune and developmental pathways. Enrichment analysis of these down-regulated genes confirmed the presence of an “ultra-immune-cold” phenotype. Highly significant GO terms included “leukocyte activation,” “regulation of leukocyte activation,” and “positive regulation of immune response,” all exhibiting exceptionally low q-values (log10(q) < -70). KEGG analysis echoed this suppression, particularly for cytokine-cytokine receptor interactions. Metascape’s cell type signature analysis further indicated a depletion of both lymphoid and myeloid lineage markers. The 15 up-regulated genes did not cluster into coherent pathways. The most enriched GO term “pattern specification process” was not statistically significant (q = 1.0), suggesting this gene set is either functionally heterogeneous or lacks defined biological convergence.

## Discussion

In this research, I utilized a multi-algorithm, unsupervised anomaly detection (AD) framework to pinpoint and analyze molecularly distinct tumor samples, termed High-Concordance Anomalies (HCAs), within the comprehensive TCGA-BRCA transcriptomic dataset. The analysis identified a consistent HCA subgroup (41 out of 1218 samples) exhibiting a complex molecular phenotype, notably characterized by significant immune suppression, specific metabolic alterations, and the atypical expression of developmental or neural-like gene programs. This study also underscores a potentially “ultra-immune-cold” and clinically significant subgroup within Basal-like breast cancer.

Employing four diverse AD algorithms and concentrating on concordantly identified HCAs provided a robust method for outlier detection, minimizing biases associated with any single algorithm. A notable finding was that 56% of these HCAs were previously classified as PAM50_Unknown based on standard RNAseq-based PAM50 classification. This suggests that the unique molecular state of HCAs, detailed herein, underlies their ambiguity to traditional classifiers. The AD concordance approach thus offers a powerful, data-driven means to identify and begin to molecularly define this challenging “dark matter” of breast cancer heterogeneity.

The predominant molecular feature of the HCA group was a pervasive “immune-cold” or “immune-exhausted” state, evidenced by the significant downregulation of nearly 1750 genes (HCAs vs. 0-vote normals). Pathway enrichment analysis (Figure 3A) confirmed this, with profound suppression of crucial immune processes such as GO:0001775: cell activation, GO:0050778: positive regulation of immune response, and hsa04060: Cytokine-cytokine receptor interaction, along with Hallmark pathways like HALLMARK INTERFERON GAMMA RESPONSE. This molecular phenotype indicates a tumor microenvironment largely deficient in effective anti-tumor immunity, a state known to impact therapeutic response, particularly to immunotherapies. While “immune-cold” tumors are a recognized concept, this study specifically links this strong, coherent signature to samples robustly identified as outliers by multiple, diverse unsupervised AD methods and highlights its strong association with PAM50_Unknown status.

**Figure 1A:**
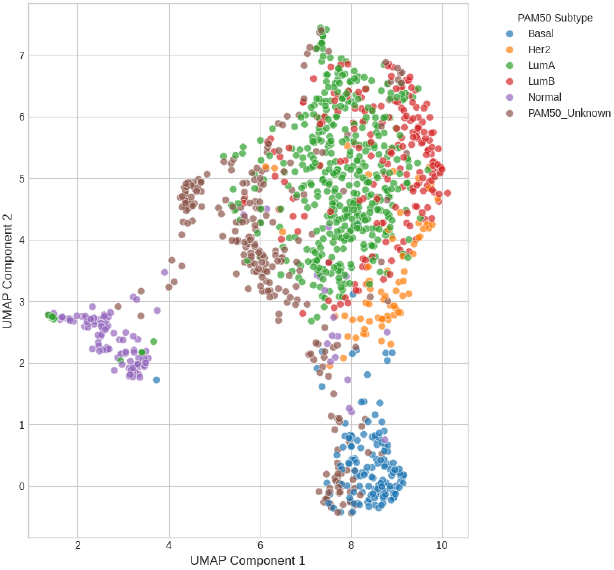
UMAP of Samples by PAM50 Subtype.

**Figure 1B:**
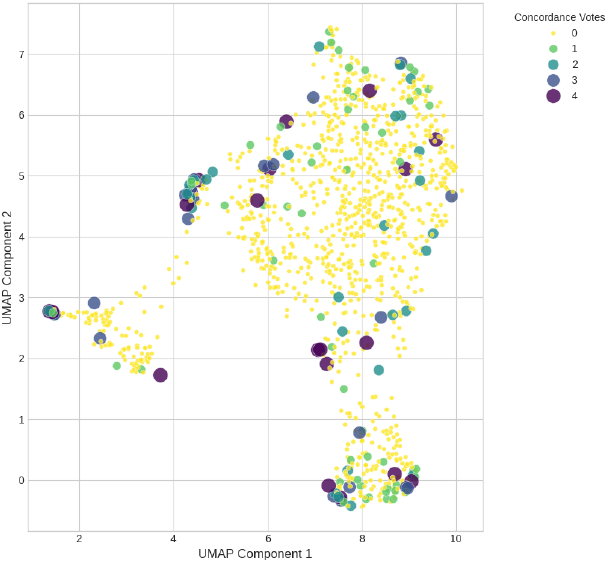
UMAP of Samples by Anomaly Concordance Votes.

**Figure 1C:**
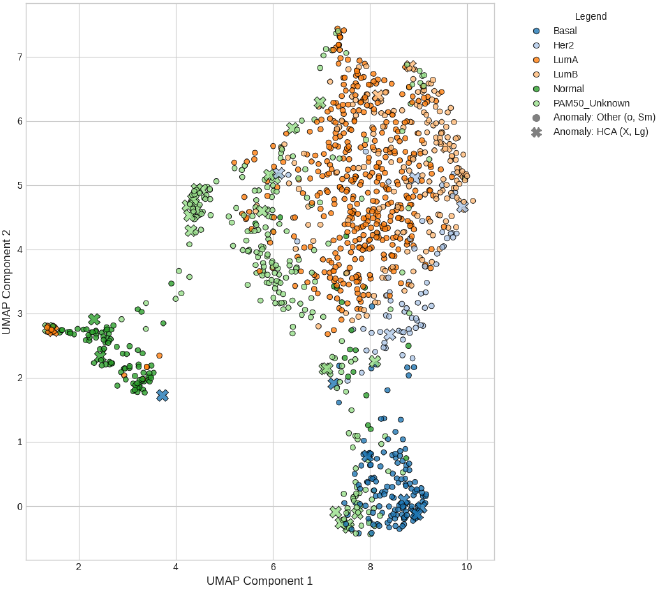
UMAP - HCAs (>2 Votes) & PAM50 Subtypes.

**Figure 2:**
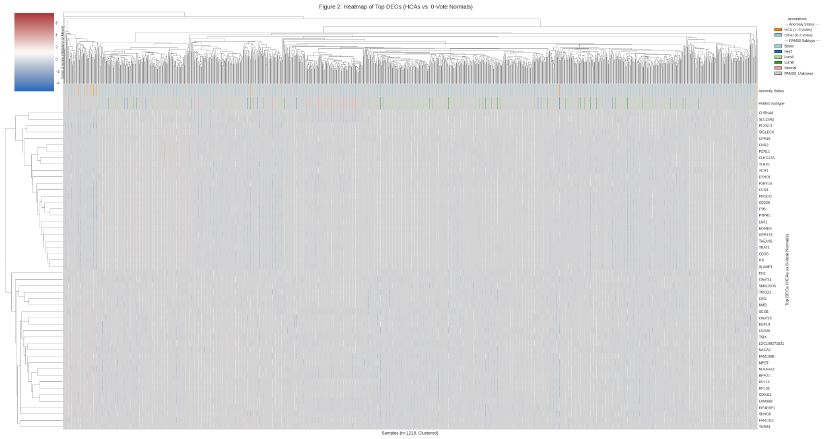

**Figure 3A:**
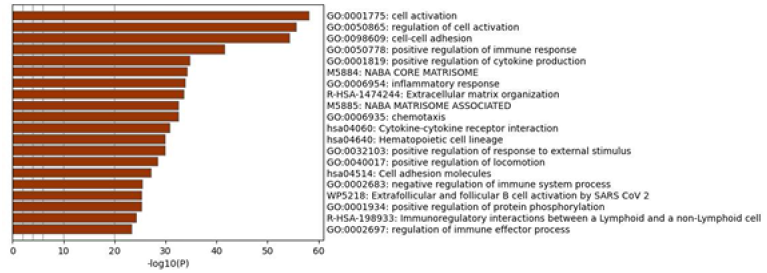

Complementing this immune quiescence, the 160 genes upregulated in HCAs (Figure 3B) suggested active cellular reprogramming. Significant enrichment for R-HSA-5660526: Response to metal ions and metabolic pathways like GO:0006091: Generation of precursor metabolites and energy indicates potential metabolic adaptations or stress responses. The intriguing upregulation of genes associated with M5883 NABA SECRETED FACTORS (ECM-related) and neural/developmental programs (e.g., GO:0007218: Neuropeptide signaling pathway) hints at possible lineage plasticity or de-differentiation, phenomena sometimes linked to tumor aggressiveness.

**Figure 3B:**
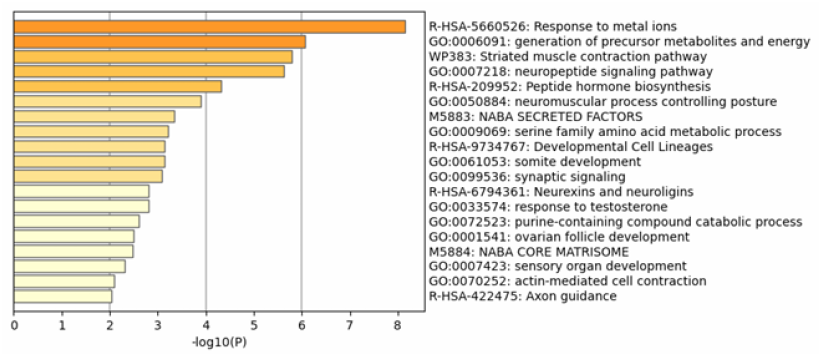

The targeted analysis of the HCA-Basal subgroup (n=7) versus “0-vote normal” Basal tumors (n=113) provided even more refined insights. These HCA-Basal samples exhibited an “ultra-immune-cold” signature, with a further 499 genes significantly downregulated compared to their typical Basal counterparts. Pathway analysis (Figure 5A) revealed an exceptionally intense suppression of GO:0045321: leukocyte activation (Log10(q) -80.33), GO:0002694: regulation of leukocyte activation (Log10(q) -71.10), and related immune pathways. The downregulation of CSF2 (GM-CSF) as a top hit in this context is notable, given its role in myeloid cell function and anti-tumor immunity. This profound immune difference within the Basal subtype may explain the visually striking trend towards poorer survival observed for the few HCA-Basal samples with OS data (n=5, p=0.1387; Figure 4B). The Basal-like subtype is notoriously heterogeneous and aggressive; the AD-derived “ultra-immune-cold” HCA-Basal signature may delineate a particularly high-risk subgroup. The 15 upregulated genes in HCA-Basal did not show strong pathway enrichment, suggesting their specific distinction from other Basals is primarily driven by this enhanced immune suppression.

**Figure 4A:**
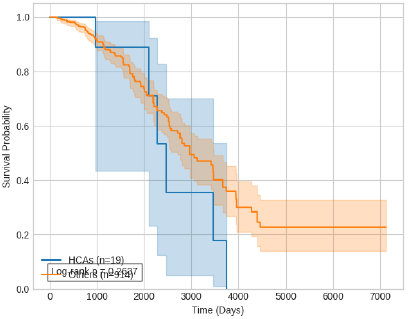

**Figure 4B:**
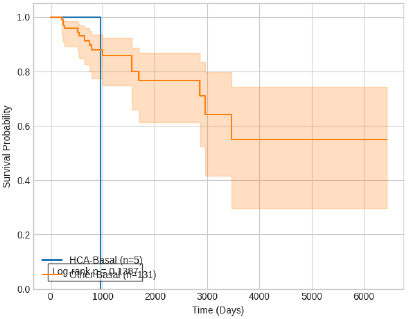

**Figure 5:**
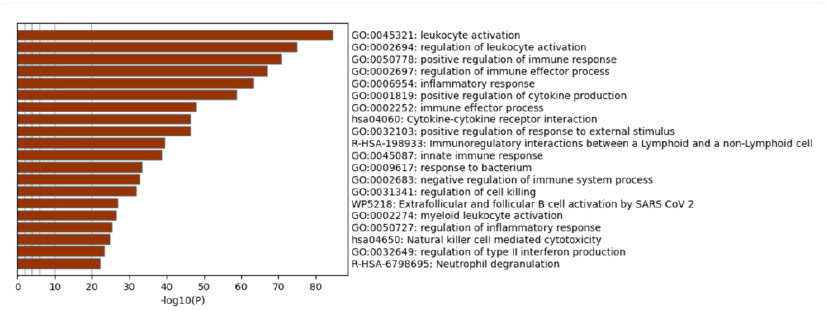

It is important to note that while the overall HCA group did not show significantly different OS (p=0.2637; Figure 4A), this likely reflects the heterogeneity within the HCAs themselves. The true clinical relevance may lie in subtype-specific contexts or in predicting response to targeted or immunotherapies. This study’s novelty arises from the specific AD concordance methodology, the comprehensive molecular definition of the HCA “immune-cold, metabolically active, developmentally divergent” phenotype, its strong link to PAM50_Unknown status, and the identification of the potentially high-risk “ultra-immune-cold” HCA-Basal variant.

## Conclusion

This study applied a multi-model unsupervised anomaly detection framework to the TCGA-BRCA transcriptomic dataset and uncovered several key insights into breast cancer heterogeneity:

A distinct group of 41 High-Concordance Anomaly (HCA) samples was identified, representing tumors with transcriptomic profiles that substantially diverge from the broader cohort, as consistently flagged by multiple anomaly detection algorithms.

These HCA samples were predominantly defined by a strongly immune-suppressed molecular signature, demonstrated by the significant downregulation of approximately 1,750 genes enriched in immune activation pathways, such as leukocyte activation and cytokine signaling.

In parallel, HCAs showed upregulation of roughly 160 genes associated with metabolic regulation, response to metal ions, and extracellular matrix organization—suggesting shifts in non-immune cellular functions and possible activation of alternative lineage or developmental programs.

Notably, over half (56%) of the HCA samples lacked definitive classification under standard PAM50 subtyping (PAM50_Unknown), suggesting that these tumors represent a biologically distinct subset inadequately captured by conventional classification methods.

Within the Basal-like subtype, a smaller group of seven HCA samples emerged as an “ultra-immune-cold” variant, marked by even stronger suppression of immune-related gene expression. Though limited by sample size, these tumors displayed a trend toward worse clinical outcomes, warranting further investigation.

In conclusion, this concordance-based anomaly detection approach highlighted a reproducible and biologically meaningful group of outlier breast tumors with distinct immunological and transcriptomic features. These findings suggest new directions for refining tumor subtyping, enhancing risk assessment, and potentially guiding therapeutic strategies aimed at poorly characterized and immune-inert tumor subpopulations.

Strengths include the rigorous multi-algorithm AD approach and the use of a large public dataset. Limitations include the retrospective nature, the current transcriptomic-only focus for AD input, small sample sizes for some subgroup analyses impacting statistical power for clinical correlations (especially survival), and the need for external experimental validation of the functional consequences of the observed molecular signatures. The “0-vote normal” baseline, while data-driven, may also contain some level of inherent tumor heterogeneity.

In conclusion, this AI-guided unsupervised AD investigation has robustly identified HCAs in TCGA-BRCA with a striking molecular phenotype. This work significantly deepens our understanding of BRCA heterogeneity, providing a molecular basis for many PAM50_Unknown tumors and identifying a potentially aggressive “ultra-immune-cold” variant within Basal-like cancer, thereby generating compelling hypotheses for future validation and therapeutic exploration.

### Future Work

The findings from this study open several important avenues for continued investigation to better understand the biology and clinical implications of the High Concordance Anomalies (HCAs) identified in breast cancer:

- **Validation in Independent Cohorts:** A critical next step is to replicate the identification of HCA samples and their associated immune-suppressed signatures in large, independent datasets such as METABRIC or publicly available GEO cohorts. This will help assess the reproducibility and generalizability of the observed patterns.
- **Signature Comparison and Literature Integration:** A comprehensive analysis comparing the HCA gene expression profiles and enriched pathways with existing breast cancer-related gene signatures including prognostic, immunological, and subtype-specific signatures—will clarify whether these anomalies reflect novel biological features or extend known categories.
- **Multi-Omics Characterization:** Integrating other layers of molecular data (e.g., somatic mutations, copy number alterations, DNA methylation, and proteomic profiles) for HCA samples will be essential to uncover potential drivers of their distinct transcriptomic state, including the severe immune suppression and a typical gene expression trends.
- **Functional Investigation of Key Genes and Pathways:** Experimental validation using in vitro or in vivo models that recapitulate the HCA phenotype would help determine the functional roles of specific differentially expressed genes such as *CSF2* in HCA-Basal and the biological impact of the dysregulated pathways.
- **Therapeutic Relevance:** Given the immune-inert nature of these tumors, further research should explore their potential resistance to immunotherapies and standard treatments. Additionally, the upregulated genes involved in metabolic regulation or metal ion response may present novel therapeutic targets specific to this subgroup.
- **Refinement of Detection Methods:** Future work should explore enhancements to the anomaly detection methodology, such as more sophisticated ensemble techniques, optimized hyperparameter tuning, and targeted feature selection strategies to better capture biologically relevant outliers.
- **Application to Other Cancer Types:** The anomaly detection framework can be extended to other cancer types within TCGA to determine whether similarly distinct transcriptomic outlier groups particularly those with immune-suppressive features exist across cancers or are unique to breast cancer.

## Supporting information

.ipynb file(jup notebook)

