## Supplementary material for "Beyond PAM50: Unsupervised Discovery of Anomalous Subgroups in Breast Cancer": .ipynb file(jup notebook): cancer_anom.ipynb

```
# %% [code]
# Cell 1: Core Imports
import numpy as np
import pandas as pd
import matplotlib.pyplot as plt
import seaborn as sns
import time
import os
!pip install lifelines
from sklearn.preprocessing import StandardScaler, MinMaxScaler
from sklearn.decomposition import PCA
from sklearn.ensemble import IsolationForest
from sklearn.svm import OneClassSVM
from sklearn.neighbors import LocalOutlierFactor
import tensorflow as tf
from tensorflow.keras.models import Model, Sequential
from tensorflow.keras.layers import Input, Dense, Dropout
from tensorflow.keras.callbacks import EarlyStopping, ReduceLROnPlateau
from tensorflow.keras.regularizers import l1_l2
import umap
from lifelines import KaplanMeierFitter
from lifelines.statistics import logrank_test
from scipy import stats
from statsmodels.stats.multitest import multipletests
# Reproducibility
np.random.seed(42)
tf.random.set_seed(42)
import pandas as pd

# %% [code]
# Cell 2: Data Loading

import pandas as pd
from google.colab import drive

drive.mount('/content/drive', force_remount=True)

base_drive_path = "/content/drive/My Drive/project/BRCA/"
file_path_expression = base_drive_path + "HiSeqV2"
file_path_clinical   = base_drive_path + "TCGA.BRCA.sampleMap_BRCA_clinicalMatrix"

df_expression_raw = pd.read_csv(file_path_expression, sep='\t', index_col=0,
                                compression='gzip' if file_path_expression.endswith('.gz') else None,
                                low_memory=False)
df_expression_transposed = df_expression_raw.transpose()
df_expression_transposed.index.name = 'sampleID'

df_clinical_raw = pd.read_csv(file_path_clinical, sep='\t', index_col=0,
                              compression='gzip' if file_path_clinical.endswith('.gz') else None,
                              low_memory=False)
df_clinical_raw.index.name = 'sampleID'

chosen_pam50_col_name = 'PAM50Call_RNAseq'

if chosen_pam50_col_name not in df_clinical_raw.columns:

    print(f"CRITICAL WARNING: Expected PAM50 column '{chosen_pam50_col_name}' not found in df_clinical_raw. Downstream analyses will be affected.")
    y_pam50_from_clinical = pd.Series(dtype='object', name=chosen_pam50_col_name)
else:
    y_pam50_from_clinical = df_clinical_raw[chosen_pam50_col_name].copy()

# %% [code]
#cell 3
import numpy as np

common_samples = df_expression_transposed.index.intersection(df_clinical_raw.index)
df_expr_aligned = df_expression_transposed.loc[common_samples]
df_clinical_aligned = df_clinical_raw.loc[common_samples]

df_expr_aligned = df_expr_aligned.apply(pd.to_numeric, errors='coerce')

if df_expr_aligned.isnull().any().any():
    df_expr_aligned = df_expr_aligned.fillna(df_expr_aligned.mean())

mean_expression_threshold = 0.5
genes_to_keep_mean = df_expr_aligned.mean(axis=0) > mean_expression_threshold
df_expr_filtered_mean = df_expr_aligned.loc[:, genes_to_keep_mean]

variance_quantile_threshold = 0.25
variances = df_expr_filtered_mean.var(axis=0)
variance_cutoff = variances.quantile(variance_quantile_threshold)
genes_to_keep_var = variances > variance_cutoff
X_processed = df_expr_filtered_mean.loc[:, genes_to_keep_var]

pam50_col_name = 'PAM50Call_RNAseq'
y_pam50 = df_clinical_aligned[pam50_col_name].copy()
y_pam50.fillna('PAM50_Unknown', inplace=True)
y_pam50.name = 'PAM50_Subtype'


os_event_col_name = 'OS_event_nature2012'
os_time_col_name = 'OS_Time_nature2012'

if os_event_col_name in df_clinical_aligned.columns and os_time_col_name in df_clinical_aligned.columns:
    y_os_event = pd.to_numeric(df_clinical_aligned[os_event_col_name], errors='coerce')
    y_os_time = pd.to_numeric(df_clinical_aligned[os_time_col_name], errors='coerce')
else:
    y_os_event = pd.Series(index=df_clinical_aligned.index, dtype='float')
    y_os_time = pd.Series(index=df_clinical_aligned.index, dtype='float')


y_os_event.name = 'OS_Event'
y_os_time.name = 'OS_Time'

age_col_name = 'age_at_initial_pathologic_diagnosis'
if age_col_name in df_clinical_aligned.columns:
    y_age = pd.to_numeric(df_clinical_aligned[age_col_name], errors='coerce')
else:
    y_age = pd.Series(index=df_clinical_aligned.index, dtype='float')
y_age.name = 'Age'


path_stage_col_name_primary = 'pathologic_stage'
path_stage_col_name_secondary = 'AJCC_Stage_nature2012'

if path_stage_col_name_primary in df_clinical_aligned.columns:
    y_stage = df_clinical_aligned[path_stage_col_name_primary].copy()
elif path_stage_col_name_secondary in df_clinical_aligned.columns:
    y_stage = df_clinical_aligned[path_stage_col_name_secondary].copy()
else:
    y_stage = pd.Series(index=df_clinical_aligned.index, dtype='object')
y_stage.name = 'Pathologic_Stage_Raw'

# %% [code]
# Cell 4: Feature Scaling & Anomaly Detection Models (Production)

#Scale Gene Expression Features
scaler_expression = StandardScaler()
X_scaled = scaler_expression.fit_transform(X_processed)
X_scaled_df = pd.DataFrame(X_scaled, index=X_processed.index, columns=X_processed.columns)

scaler_ae = MinMaxScaler()
X_scaled_ae = scaler_ae.fit_transform(X_processed)
X_scaled_ae_df = pd.DataFrame(X_scaled_ae, index=X_processed.index, columns=X_processed.columns)

# Define and Apply Anomaly Detection Models
contamination_rate = 0.05
anomaly_predictions = pd.DataFrame(index=X_processed.index)

# A. Isolation Forest
if_model = IsolationForest(n_estimators=200, contamination=contamination_rate,
                           max_samples='auto', random_state=42, n_jobs=-1)
if_pred = if_model.fit_predict(X_scaled_df)
anomaly_predictions['IsolationForest'] = (if_pred == -1).astype(int)

# B. One-Class SVM
ocsvm_model = OneClassSVM(kernel='rbf', gamma='auto', nu=contamination_rate)
ocsvm_pred = ocsvm_model.fit_predict(X_scaled_df)
anomaly_predictions['OneClassSVM'] = (ocsvm_pred == -1).astype(int)

# C. Local Outlier Factor (LOF)
lof_model = LocalOutlierFactor(n_neighbors=30, contamination=contamination_rate,
                               novelty=False, n_jobs=-1)
lof_pred = lof_model.fit_predict(X_scaled_df)
anomaly_predictions['LOF'] = (lof_pred == -1).astype(int)

# D. Autoencoder
input_dim_ae = X_scaled_ae_df.shape[1]
encoding_dim_l1 = max(64, int(input_dim_ae / 20))
encoding_dim_l2 = max(32, int(encoding_dim_l1 / 2))
bottleneck_dim = max(16, int(encoding_dim_l2 / 2))

autoencoder = Sequential([
    Input(shape=(input_dim_ae,)),
    Dense(encoding_dim_l1, activation='relu', kernel_regularizer=l1_l2(l1=1e-5, l2=1e-5)),
    Dropout(0.1),
    Dense(encoding_dim_l2, activation='relu', kernel_regularizer=l1_l2(l1=1e-5, l2=1e-5)),
    Dropout(0.1),
    Dense(bottleneck_dim, activation='relu', name='bottleneck_layer', kernel_regularizer=l1_l2(l1=1e-5, l2=1e-5)),
    Dense(encoding_dim_l2, activation='relu', kernel_regularizer=l1_l2(l1=1e-5, l2=1e-5)),
    Dropout(0.1),
    Dense(encoding_dim_l1, activation='relu', kernel_regularizer=l1_l2(l1=1e-5, l2=1e-5)),
    Dropout(0.1),
    Dense(input_dim_ae, activation='sigmoid')
])
autoencoder.compile(optimizer=tf.keras.optimizers.Adam(learning_rate=0.001), loss='mse')

early_stopping_ae = EarlyStopping(monitor='val_loss', patience=15, restore_best_weights=True, verbose=0)
reduce_lr_ae = ReduceLROnPlateau(monitor='val_loss', factor=0.2, patience=7, min_lr=1e-6, verbose=0)

history = autoencoder.fit(X_scaled_ae_df, X_scaled_ae_df,
                          epochs=200, batch_size=32, shuffle=True,
                          validation_split=0.15,
                          callbacks=[early_stopping_ae, reduce_lr_ae],
                          verbose=0)

X_pred_ae = autoencoder.predict(X_scaled_ae_df, verbose=0)
mse_ae = np.mean(np.power(X_scaled_ae_df - X_pred_ae, 2), axis=1)
reconstruction_errors_ae = pd.Series(mse_ae, index=X_scaled_ae_df.index)
ae_error_threshold = np.percentile(reconstruction_errors_ae, 100 * (1 - contamination_rate))
anomaly_predictions['Autoencoder'] = (reconstruction_errors_ae > ae_error_threshold).astype(int)

anomaly_predictions['Concordance_Votes'] = anomaly_predictions.sum(axis=1)


# %% [code]
# Cell 5: UMAP Visualization & Core Anomaly Characterization Data

import pandas as pd
import numpy as np
import matplotlib.pyplot as plt
import seaborn as sns
import umap
from scipy import stats


reducer_for_viz = umap.UMAP(n_neighbors=25, min_dist=0.1, n_components=2,
                            metric='euclidean', random_state=42, n_jobs=1)
X_umap_coords = reducer_for_viz.fit_transform(X_scaled_df)
df_umap_coords = pd.DataFrame(X_umap_coords, columns=['UMAP1', 'UMAP2'], index=X_scaled_df.index)


df_viz = pd.concat([
    df_umap_coords,
    anomaly_predictions[['IsolationForest', 'OneClassSVM', 'LOF', 'Autoencoder', 'Concordance_Votes']],
    y_pam50
], axis=1)

if 'PAM50_Subtype' not in df_viz.columns and y_pam50.name in df_viz.columns:
    df_viz.rename(columns={y_pam50.name: 'PAM50_Subtype'}, inplace=True)
elif 'PAM50_Subtype' not in df_viz.columns and y_pam50.name not in df_viz.columns and 'PAM50Call_RNAseq' in df_viz.columns:
    df_viz.rename(columns={'PAM50Call_RNAseq': 'PAM50_Subtype'}, inplace=True)

#Generate UMAP Visualizations
plt.style.use('seaborn-v0_8-whitegrid')
min_votes_for_high_concordance = 3
pam50_order_for_plots = sorted(df_viz['PAM50_Subtype'].unique())

#UMAP by PAM50 Subtype
plt.figure(figsize=(10, 8))
sns.scatterplot(x='UMAP1', y='UMAP2', hue='PAM50_Subtype', hue_order=pam50_order_for_plots,
                palette='tab10', data=df_viz, legend='full', s=50, alpha=0.7)
plt.title('Figure 1A: UMAP of Samples by PAM50 Subtype', fontsize=14)
plt.xlabel('UMAP Component 1'); plt.ylabel('UMAP Component 2')
plt.legend(title='PAM50 Subtype', bbox_to_anchor=(1.05, 1), loc='upper left')
plt.tight_layout(rect=[0,0,0.85,1]); plt.show()

#UMAP by Anomaly Concordance Votes
plt.figure(figsize=(10, 8))
sns.scatterplot(x='UMAP1', y='UMAP2', hue='Concordance_Votes', size='Concordance_Votes',
                sizes=(20, 200), palette='viridis_r', data=df_viz, legend='full', alpha=0.8)
plt.title('Figure 1B: UMAP by Anomaly Concordance Votes', fontsize=14)
plt.xlabel('UMAP Component 1'); plt.ylabel('UMAP Component 2')
plt.legend(title='Concordance Votes', bbox_to_anchor=(1.05, 1), loc='upper left')
plt.tight_layout(rect=[0,0,0.85,1]); plt.show()

#UMAP Highlighting HCAs with PAM50 Subtypes
df_viz['High_Concordance_Anomaly'] = (df_viz['Concordance_Votes'] >= min_votes_for_high_concordance)
plt.figure(figsize=(12, 9))
scatter_plot_1c = sns.scatterplot(
    x='UMAP1', y='UMAP2', hue='PAM50_Subtype', hue_order=pam50_order_for_plots,
    size='High_Concordance_Anomaly', sizes={True: 150, False: 30},
    style='High_Concordance_Anomaly', markers={True: 'X', False: 'o'},
    palette='tab20', data=df_viz, alpha=0.8, edgecolor='black', linewidth=0.7)
plt.title(f'Figure 1C: UMAP - HCAs (Votes >={min_votes_for_high_concordance}) & PAM50', fontsize=14)
plt.xlabel('UMAP Component 1'); plt.ylabel('UMAP Component 2')

handles_1c, labels_1c = scatter_plot_1c.get_legend_handles_labels()
unique_legend_map_1c = {label: handle for handle, label in reversed(list(zip(handles_1c, labels_1c)))}
plt.legend(unique_legend_map_1c.values(), unique_legend_map_1c.keys(), title='Legend', bbox_to_anchor=(1.05, 1), loc='upper left', fontsize=9)
plt.tight_layout(rect=[0,0,0.82,1]); plt.show()

normal_samples_in_viz = df_viz[~df_viz['High_Concordance_Anomaly']]


df_viz_with_age = df_viz.merge(y_age.rename('Age'), left_index=True, right_index=True, how='left')
age_anomalies_data = df_viz_with_age[df_viz_with_age['High_Concordance_Anomaly']]['Age'].dropna()
age_normals_data = df_viz_with_age[~df_viz_with_age['High_Concordance_Anomaly']]['Age'].dropna()

if 'Age' in df_viz_with_age.columns and not age_anomalies_data.empty and not age_normals_data.empty:
    plt.figure(figsize=(6, 7))
    sns.boxplot(x='High_Concordance_Anomaly', y='Age', hue='High_Concordance_Anomaly',
                data=df_viz_with_age, palette=['royalblue', 'orangered'], legend=False)
    plt.title('Age Distribution by Anomaly Status', fontsize=14)
    plt.xticks(ticks=[0, 1], labels=['Other (0-2 Votes)', f'HCA (Votes >={min_votes_for_high_concordance})'])
    plt.ylabel('Age at Diagnosis'); plt.xlabel('Anomaly Status')
    plt.tight_layout(); plt.show()


# %% [code]
# Cell 6: Molecular (DGE) & Clinical (Survival) Characterization (Production)

import pandas as pd
import numpy as np
import matplotlib.pyplot as plt
from scipy import stats
from statsmodels.stats.multitest import multipletests
from lifelines import KaplanMeierFitter
from lifelines.statistics import logrank_test


#Isolate Anomaly Groups and Baseline Normals ---
min_votes_for_high_concordance = 3
is_high_concordance_anomaly = (anomaly_predictions['Concordance_Votes'] >= min_votes_for_high_concordance)
high_concordance_indices = X_processed.index[is_high_concordance_anomaly]
X_anomalies_hc = X_processed.loc[high_concordance_indices]
normal_0_votes_indices = X_processed.index[anomaly_predictions['Concordance_Votes'] == 0]
X_normals_0v = X_processed.loc[normal_0_votes_indices]

#Differential Gene Expression (Global HCAs vs. 0-Vote Normals)
df_dge = pd.DataFrame()
sig_genes_up = pd.DataFrame()
sig_genes_down = pd.DataFrame()

if len(X_anomalies_hc) >= 2 and len(X_normals_0v) >= 2:
    dge_results_list = []
    for gene in X_processed.columns:
        group1_expr = X_anomalies_hc[gene].dropna()
        group2_expr = X_normals_0v[gene].dropna()
        if len(group1_expr) < 2 or len(group2_expr) < 2: continue
        log2_fold_change = group1_expr.mean() - group2_expr.mean()
        t_stat, p_value = stats.ttest_ind(group1_expr, group2_expr, equal_var=False, nan_policy='omit')
        dge_results_list.append({'gene': gene, 'log2FC': log2_fold_change, 'p_value': p_value, 't_stat': t_stat})

    if dge_results_list:
        df_dge = pd.DataFrame(dge_results_list)
        if not df_dge.empty:
            df_dge.dropna(subset=['p_value'], inplace=True)
            if not df_dge.empty:
                reject, q_values, _, _ = multipletests(df_dge['p_value'], method='fdr_bh')
                df_dge['q_value'] = q_values
                # df_dge['significant_bh'] = reject
                df_dge_sorted = df_dge.sort_values(by='q_value')
                sig_genes_up = df_dge_sorted[(df_dge_sorted['q_value'] < 0.05) & (df_dge_sorted['log2FC'] > 1.0)]
                sig_genes_down = df_dge_sorted[(df_dge_sorted['q_value'] < 0.05) & (df_dge_sorted['log2FC'] < -1.0)]

#Save Gene Lists for Pathway Analysis (Global HCAs)
if not sig_genes_up.empty:
    sig_genes_up[['gene', 'log2FC', 'p_value', 'q_value']].to_csv('significant_UP_genes_anomalies_vs_0vote.tsv', sep='\t', index=False)
if not sig_genes_down.empty:
    sig_genes_down[['gene', 'log2FC', 'p_value', 'q_value']].to_csv('significant_DOWN_genes_anomalies_vs_0vote.tsv', sep='\t', index=False)
if 'df_dge_sorted' in globals() and not df_dge_sorted.empty:
    df_dge_sorted[['gene', 'log2FC', 'p_value', 'q_value']].to_csv('ALL_genes_ranked_DGE_anomalies_vs_0vote.tsv', sep='\t', index=False)

#Survival Analysis
T_aligned = pd.to_numeric(y_os_time.loc[X_processed.index], errors='coerce').rename('Time')
E_aligned = pd.to_numeric(y_os_event.loc[X_processed.index], errors='coerce').rename('Event')

#HCAs vs. Other Samples
df_survival_hc_vs_other = pd.concat([T_aligned, E_aligned, is_high_concordance_anomaly.rename('Is_Anomaly_HC')], axis=1)
df_survival_hc_vs_other.dropna(subset=['Time', 'Event'], inplace=True)

if len(df_survival_hc_vs_other['Is_Anomaly_HC'].value_counts()) >= 2 and df_survival_hc_vs_other['Is_Anomaly_HC'].value_counts().min() >= 2:
    kmf_plotter_A = KaplanMeierFitter()
    plt.figure(figsize=(8, 6))
    ax_km_A = None
    group1_A = df_survival_hc_vs_other[df_survival_hc_vs_other['Is_Anomaly_HC']]
    group2_A = df_survival_hc_vs_other[~df_survival_hc_vs_other['Is_Anomaly_HC']]
    if len(group1_A) > 1: kmf_plotter_A.fit(group1_A['Time'], group1_A['Event'], label=f'HCAs (n={len(group1_A)})'); ax_km_A = kmf_plotter_A.plot_survival_function()
    if len(group2_A) > 1: kmf_plotter_A.fit(group2_A['Time'], group2_A['Event'], label=f'Others (0-2 Votes) (n={len(group2_A)})'); kmf_plotter_A.plot_survival_function(ax=ax_km_A if ax_km_A else None)

    if ax_km_A:
        plt.title('Figure 4A: OS - HCAs vs. Others'); plt.xlabel('Time (Days)'); plt.ylabel('Survival Probability'); plt.grid(True); plt.ylim(0,1.05)
        if len(group1_A) > 1 and len(group2_A) > 1:
            res_A = logrank_test(group1_A['Time'], group2_A['Time'], group1_A['Event'], group2_A['Event'])
            plt.text(0.05, 0.05, f'Log-rank p = {res_A.p_value:.4f}', transform=ax_km_A.transAxes, bbox=dict(facecolor='white', alpha=0.7))
        plt.show()

# B. HCA-Basal vs. Other-Basal
df_survival_pam50 = pd.concat([T_aligned, E_aligned, y_pam50.rename('PAM50_Subtype'), is_high_concordance_anomaly.rename('Is_Anomaly_HC')], axis=1)
df_survival_pam50.dropna(subset=['Time', 'Event', 'PAM50_Subtype'], inplace=True)
df_survival_pam50 = df_survival_pam50[df_survival_pam50['PAM50_Subtype'] != 'PAM50_Unknown']

if 'Basal' in df_survival_pam50['PAM50_Subtype'].unique():
    df_basal_survival = df_survival_pam50[df_survival_pam50['PAM50_Subtype'] == 'Basal']
    if 'Is_Anomaly_HC' in df_basal_survival.columns and len(df_basal_survival['Is_Anomaly_HC'].value_counts()) >= 2 and df_basal_survival['Is_Anomaly_HC'].value_counts().min() >= 2 :
        kmf_plotter_B = KaplanMeierFitter()
        plt.figure(figsize=(8,6))
        ax_km_B = None
        group1_B = df_basal_survival[df_basal_survival['Is_Anomaly_HC']]
        group2_B = df_basal_survival[~df_basal_survival['Is_Anomaly_HC']]
        if len(group1_B) > 1: kmf_plotter_B.fit(group1_B['Time'], group1_B['Event'], label=f'HCA-Basal (n={len(group1_B)})'); ax_km_B = kmf_plotter_B.plot_survival_function()
        if len(group2_B) > 1: kmf_plotter_B.fit(group2_B['Time'], group2_B['Event'], label=f'Other Basal (n={len(group2_B)})'); kmf_plotter_B.plot_survival_function(ax=ax_km_B if ax_km_B else None)

        if ax_km_B:
            plt.title('Figure 4B: OS - HCA-Basal vs. Other Basal'); plt.xlabel('Time (Days)'); plt.ylabel('Survival Probability'); plt.grid(True); plt.ylim(0,1.05)
            if len(group1_B) > 1 and len(group2_B) > 1:
                res_B = logrank_test(group1_B['Time'], group2_B['Time'], group1_B['Event'], group2_B['Event'])
                plt.text(0.05, 0.05, f'Log-rank p = {res_B.p_value:.4f}', transform=ax_km_B.transAxes, bbox=dict(facecolor='white', alpha=0.7))
            plt.show()


# %% [code]
# Cell 8: Heatmap Visualization of Top DEGs
import matplotlib.pyplot as plt
import seaborn as sns

# Select Top DEGs
n_top_genes = 25
sig_genes_from_dge = df_dge_sorted[
    (df_dge_sorted['q_value'] < 0.05) & (np.abs(df_dge_sorted['log2FC']) > 1.0)
]
top_up_genes = sig_genes_from_dge[sig_genes_from_dge['log2FC'] > 0].nsmallest(n_top_genes, 'q_value')['gene'].tolist()
top_down_genes = sig_genes_from_dge[sig_genes_from_dge['log2FC'] < 0].nsmallest(n_top_genes, 'q_value')['gene'].tolist()
selected_genes = sorted(list(set(top_up_genes + top_down_genes)))

if selected_genes:
    #Prepare Data for Heatmap
    valid_selected_genes = [gene for gene in selected_genes if gene in X_processed.columns]
    if valid_selected_genes:
        X_heatmap_data = X_processed[valid_selected_genes]

        #Create Sample Annotations for Heatmap Columns
        anomaly_status_labels = is_high_concordance_anomaly.loc[X_heatmap_data.index].rename("Anomaly_Status_Labels").map({True: 'HCA (>=3 Votes)', False: 'Other (0-2 Votes)'})
        anomaly_status_color_map = {'HCA (>=3 Votes)': 'darkorange', 'Other (0-2 Votes)': 'skyblue'}

        pam50_subtype_labels = y_pam50.loc[X_heatmap_data.index].rename("PAM50_Subtype_Labels")
        unique_pam50_values = sorted(pam50_subtype_labels.unique())
        pam50_base_palette = sns.color_palette("Paired", n_colors=max(12, len(unique_pam50_values)))
        pam50_color_map = {subtype: pam50_base_palette[i % len(pam50_base_palette)] for i, subtype in enumerate(unique_pam50_values)}
        if 'PAM50_Unknown' in pam50_color_map: pam50_color_map['PAM50_Unknown'] = 'lightgrey'

        df_col_colors_for_clustermap = pd.DataFrame(index=X_heatmap_data.index)
        df_col_colors_for_clustermap['Anomaly Status'] = anomaly_status_labels.map(anomaly_status_color_map)
        df_col_colors_for_clustermap['PAM50 Subtype'] = pam50_subtype_labels.map(pam50_color_map)

        #Generate Heatmap using seaborn.clustermap
        X_heatmap_plot_data = X_heatmap_data.transpose()

        if not X_heatmap_plot_data.empty:
            if not X_heatmap_plot_data.apply(lambda s: pd.to_numeric(s, errors='coerce').notnull().all()).all():
                X_heatmap_plot_data = X_heatmap_plot_data.apply(pd.to_numeric, errors='coerce').fillna(0)

            plot_height = np.clip(len(selected_genes) / 3.0, 10, 30)
            plot_width = np.clip(X_heatmap_plot_data.shape[1] / 40.0, 12, 35)

            g = sns.clustermap(X_heatmap_plot_data,
                               z_score=0, cmap="vlag", col_colors=df_col_colors_for_clustermap,
                               linewidths=0.1, linecolor='lightgray', figsize=(plot_width, plot_height),
                               metric="euclidean", method="average",
                               yticklabels=True, xticklabels=False,
                               dendrogram_ratio=(.08, .18),
                               cbar_kws={'label': 'Gene Expression Z-score'})

            g.ax_heatmap.set_ylabel("Top DEGs", fontsize=12)
            g.ax_heatmap.set_xlabel(f"Samples (n={X_heatmap_plot_data.shape[1]}, Clustered)", fontsize=12)
            plt.suptitle("Figure 2: Heatmap of Top DEGs (HCAs vs. 0-Vote Normals)", y=1.00, fontsize=16)

            from matplotlib.patches import Patch
            legend_elements = []
            legend_elements.append(Patch(facecolor='white', edgecolor='white', label='Anomaly Status:'))
            for label, color in anomaly_status_color_map.items():
                legend_elements.append(Patch(facecolor=color, edgecolor='black', label=label))
            legend_elements.append(Patch(facecolor='white', edgecolor='white', label='PAM50 Subtype:'))
            for label, color in sorted(pam50_color_map.items()):
                legend_elements.append(Patch(facecolor=color, edgecolor='black', label=label))

            if hasattr(g, 'ax_col_dendrogram') and g.ax_col_dendrogram:
                 g.ax_col_dendrogram.legend(handles=legend_elements, loc="center left",
                                           bbox_to_anchor=(1.02, 0.5),
                                           title="Annotations", frameon=False)
            plt.show()

# %% [code]
# Cell 9: Targeted DGE (HCA-Basal vs. 0-Vote-Normal-Basal) (Production)

from statsmodels.stats.multitest import multipletests


# Identify Sample Subsets for Targeted DGE
min_votes_for_high_concordance = 3
is_hca = (anomaly_predictions['Concordance_Votes'] >= min_votes_for_high_concordance)
is_basal = (y_pam50 == 'Basal')
is_0_vote_normal = (anomaly_predictions['Concordance_Votes'] == 0)

hca_basal_indices = X_processed.index[is_hca & is_basal]
X_hca_basal = X_processed.loc[hca_basal_indices]

normal_0v_basal_indices = X_processed.index[is_0_vote_normal & is_basal]
X_normal_0v_basal = X_processed.loc[normal_0v_basal_indices]

#Perform Differential Gene Expression

df_dge_basal_sorted = pd.DataFrame()
sig_genes_up_basal = pd.DataFrame()
sig_genes_down_basal = pd.DataFrame()

if len(X_hca_basal) >= 2 and len(X_normal_0v_basal) >= 2:
    dge_results_basal_list = []
    for gene in X_processed.columns:
        group1_expr = X_hca_basal[gene].dropna()
        group2_expr = X_normal_0v_basal[gene].dropna()
        if len(group1_expr) < 2 or len(group2_expr) < 2: continue
        log2_fold_change = group1_expr.mean() - group2_expr.mean()
        t_stat, p_value = stats.ttest_ind(group1_expr, group2_expr, equal_var=False, nan_policy='omit')
        dge_results_basal_list.append({'gene': gene, 'log2FC': log2_fold_change, 'p_value': p_value, 't_stat': t_stat})

    if dge_results_basal_list:
        df_dge_basal = pd.DataFrame(dge_results_basal_list)
        if not df_dge_basal.empty:
            df_dge_basal.dropna(subset=['p_value'], inplace=True)
            if not df_dge_basal.empty:
                reject_basal, q_values_basal, _, _ = multipletests(df_dge_basal['p_value'], method='fdr_bh')
                df_dge_basal['q_value'] = q_values_basal
                df_dge_basal_sorted = df_dge_basal.sort_values(by='q_value')

                sig_genes_up_basal = df_dge_basal_sorted[(df_dge_basal_sorted['q_value'] < 0.05) & (df_dge_basal_sorted['log2FC'] > 1.0)]
                sig_genes_down_basal = df_dge_basal_sorted[(df_dge_basal_sorted['q_value'] < 0.05) & (df_dge_basal_sorted['log2FC'] < -1.0)]

# Save Gene Lists for Pathway Analysis
if not sig_genes_up_basal.empty:
    sig_genes_up_basal[['gene', 'log2FC', 'p_value', 'q_value']].to_csv('significant_UP_genes_HCA_Basal_vs_Normal_Basal.tsv', sep='\t', index=False)
if not sig_genes_down_basal.empty:
    sig_genes_down_basal[['gene', 'log2FC', 'p_value', 'q_value']].to_csv('significant_DOWN_genes_HCA_Basal_vs_Normal_Basal.tsv', sep='\t', index=False)
if not df_dge_basal_sorted.empty:
    df_dge_basal_sorted[['gene', 'log2FC', 'p_value', 'q_value']].to_csv('ALL_genes_ranked_DGE_HCA_Basal_vs_Normal_Basal.tsv', sep='\t', index=False)
```
